# Replication elongates short DNA, reduces sequence bias, and develops trimer structure

**DOI:** 10.1101/2023.04.28.538682

**Authors:** Adriana Calaça Serrão, Felix T. Dänekamp, Zsófia Meggyesi, Dieter Braun

## Abstract

The origin of molecular evolution required the replication of short oligonucleotides to form longer polymers. Prebiotically plausible oligonucleotide pools tend to contain more of some nucleotide bases than others. It has been unclear whether this initial bias persists and how it affects replication. To investigate this, we examined the evolution of 12 mer biased short DNA pools during enzymatic templated polymerization. Our analysis using next-generation sequencing from different time points revealed that the initial nucleotide bias of the pool disappeared in the elongated pool after isothermal replication. In contrast, the nucleotide composition at each position in the elongated sequences remained biased and varied with both position and initial bias. Furthermore, we observed the emergence of highly periodic dimer and trimer motifs in the rapidly elongated sequences. This shift in nucleotide composition and the emergence of structure through templated replication could help explain how biased prebiotic pools could undergo molecular evolution and lead to complex functional nucleic acids.

## Introduction

The replication of short oligonucleotides to create longer polymers is a central step in the origin of more functional nucleic acids. It has been addressed through enzymatic [1, 2] and non-enzymatic replication [3–5], mostly from specific sequences or naive pools of short oligomers. However, condensation of single base mononucleotides in a primordial context often leads to short oligomer pools with a sequence bias, namely with one nucleobase incorporated more into the product strands [6–9]. This bias, on the one hand, may be due to an imbalanced abundance in the environment caused by different rates of nucleotide formation and degradation in different conditions [10–14]. On the other hand, even when the environment has equimolar concentrations of all reacting nucleobases, the rate of nucleotide condensation reactions themselves may also vary for different nucleotides [6, 9, 15].

Functional nucleic acid strands are usually long, with several tens or hundreds of base pairs [16], and have specific secondary structure [17, 18]. Even though such catalytic nucleic acids occupy only a subsection of the possible sequence space [19, 20], they are still more compositionally diverse than the biased pools obtained from nucleotide condensation studies [10]. The mechanism through which such functional strands evolve from a pool of short biased oligomers, both elongating and driving the evolution of sequence information, is not fully understood [21, 22].

Templated replication is a potential mechanism through which both the compositional diversity and sequence length can increase to facilitate the exploration of sequence space while replicating sequence information [10]. Due to the complementarity of Watson-Crick base-pairing, necessary for templation, a strong bias to one nucleobase leads to the complementary base being correspondingly more incorporated in the nascent strand. This in turn homogenizes the average pool nucleotide fraction, Figure 1. While the overall nucleotide composition is expected to diversify, several studies have shown that templated replication can act as a selection mechanism in itself, enriching specific sequence motifs [1, 23–25]. More experimental investigations are needed to grasp the influence of the initial biases of the pool on the sequence level. The goal of our study was to specifically understand which motifs are enriched starting from such biased initial pools and whether the replicated pool holds memory of the initial bias.

**Figure 1:**
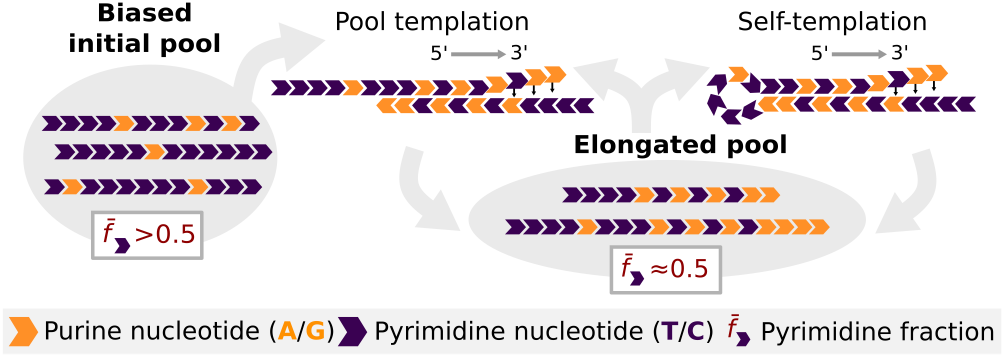
Polymerization starts from a binary initial pool (AT or GC) with a bias 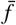 of either purine (orange) or pyrimidine (purple) nucleotides. Distinct sequences from the pool base-pair to form short duplexes and are enzymatically extended (5′-3′) complementarily to the template (“pool templation”). Longer sequences may also self-template through hairpin-like secondary structures (“self-templation”). The biased pyrimidine fraction 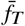 or 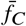 in the initial pool is countered by complementary elongation.

In contemporary biology, strand separation and elongation occur in tandem [26, 27]. The displacement of any pre-hybridized strands is performed enzymatically. However, strand displacement can also be triggered by the hybridization of other sequences in the pool [28]. This non-enzymatic strand displacement has recently been described for a prebiotic RNA replication system [29]. When compared to other prebiotic mechanisms proposed for strand separation, such as pH [22, 30], heat and salt fluctuations [31], strand displacement has the advantage that it can also occur isothermally and with a constant chemical environment [28]. It thereby erases the need for a specific set of cycle conditions that are potentially more difficult to satisfy and isolates the impact of replication on sequence structure from other environment variables.

We investigated how the sequence landscape of short biased DNA pools evolves upon templated polymerization with *Bacillus stearothermophilus* strand displacing polymerase (*Bst*). *Bst* binds to double stranded regions and elongates the strand in the 5′ to 3′ direction with high-fidelity [32–34], displacing downstream bound strands, Supplementary Information (SI) section II. A single strand can therefore go through several replication rounds, even in isothermal conditions – first through pool templation and later, when a certain length threshold is crossed, through self-templation, Figure 1. We started with a pool of short 12 mer DNA strands, with a binary composition of either AT or GC, and of all the four possible biases (A-rich, G-rich, etc.).

After following the sequence space over the course of incubation with *Bst*, we found that the initial nucleotide bias of the pool disappears, so that the resulting pool has a nucleotide fraction of 0.5 (i.e. 50%A and 50%T). While this new pool is now homogenized in terms of overall nucleotide composition, individual segments of the elongated strands still retain traces of the initial bias, due to the directionality of the polymerization. This shows that even though the overall pool nucleotide fraction changes through replication, the structure within sequences depends on the initial state. Furthermore, we have also observed that highly periodic motifs are present in sequences that elongate fast.

## Results and Discussion

Our starting pools with 10μm total DNA were composed of random 12 nucleotide (nt) long single stranded binary sequences (AT or GC only) with a bias in the nucleotide fraction, see methods section. The sequence space was thus 212 = 4096, but sequences were not represented equally due to the bias. From these initial pools, sequences were isothermally amplified with the strand-displacing enzyme *Bst*. The incubation temperatures were 35°C for AT pools and 65°C for GC pools. In a temperature screening, these led to the most extensive elongation, SI section VII.

The evolution of sequence lengths over time was analyzed through polyacrylamide gel electrophoresis (PAGE), Figure **2 a** and **b**. Different time points were analyzed for AT and GC pools to account for the different kinetics of nucleotide incorporation, Figure 2 **c** and **d**. The polymerization was stopped at 32h for AT and 8h for GC when the length distribution reached its steady state in the PAGE gel. Both the A-biased (A_0_) and the G-biased (G_0_) pools displayed replication to sequences longer than 100 mer within the first 2h. In case of the A_0_ pool, most of the short initial sequences (< 20nt) were depleted after 2h, whereas for G_0_, these remained detectable even for later time points. The remaining pools, T-biased (T_0_) and C-biased (C_0_), exhibit similar length distribution kinetics, SI section V.

**Figure 2:**
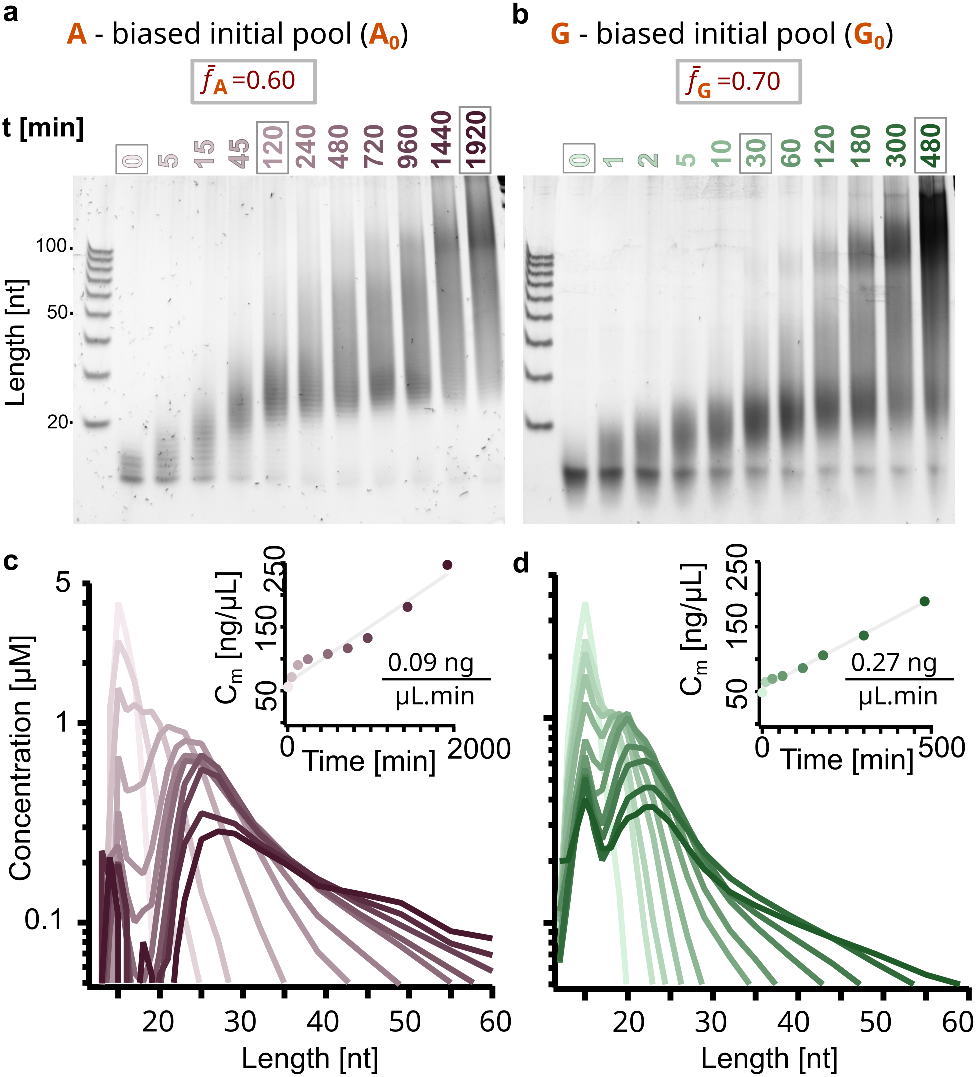
Templated polymerization of random DNA 12 mers leads to products longer than 100 mer. PAGE analysis shows the length distribution of **a** A-biased (A_0_) and **b** G-biased (G_0_) pools over time. The molar concentration of sequences was quantified and plotted over sequence length for each time point corresponding to individual lanes. A_0_ corresponds to pink (**c**) and G_0_ to green (**d**), with hue increasing over time. The total DNA mass concentration grows linearly with time (inlets) and was fitted in grey. The concentrations were obtained from the gels by PAGE smear quantification, SI section VIII and IX.

The concentration profiles over strand length were obtained via ladder-calibrated SYBR Gold fluorescence intensity in PAGE gels and depicted for all time points in Figure 2 **c** and **d**, SI sections VIII and IX. For both A_0_ and G_0_ pools, the molar concentration at later time points forms a double peaked length distribution with a long tail which continues to lengths longer than 300 nt. The first peak, around 12 nt, could be explained by the sequences of the initial pool that were not recruited for replication. The second peak, between 20 and 30 nt, could be due to fully hybridized duplexes that have a melting temperature above the incubation temperature [35].

While the total number of sequences is constant because single nucleotides get added to already existing sequences, the total DNA mass increases linearly with time as more nucleotides are incorporated, Figure 2 **c** and **d**, inlets. The difference in kinetics observed (about three times slower for AT experiments) can be explained both by the temperature dependent efficiency of *Bst* and nucleotide-dependent differences in the rate of incorporation [36, 37].

To assess the sequence content of our product strands we used Next-Generation Sequencing (NGS). For each of the four initial pools three timepoint samples were sequenced (indicated in Figure 2 **a** and **b** by the grey outlines). Respectively, these represent the initial pool, an early time point, from which we learned about fast replicating sequences, and a late time point to understand close-to-equilibrium sequence distribution. In the case of A_0_ and T_0_, we sequenced the samples at 0h, 2h and 32h, whereas in the case of G_0_ and C_0_, at 0h, 0.5h and 8h. The resulting data sets allowed us to characterize how the bias in the initial pools affects the pool evolution on short and long time scales.

The analysis of the AT data sets (A_0_ and T_0_) is depicted in Figure 3. As polymerization leads to all possible integer lengths from the initial 12 mer sequences (to the maximum range), we plotted the fraction *f*_*T*_^(*i*)^ of nucleotide T at each position *i* for sequences of the same length. We then stacked the graphs so that the positions align across lengths, Figure 3 **a, b**. The position is plotted in the direction 5′ to 3′ end, the same direction as *Bst* elongates the sequences. This way, for every sequence length, the probability of finding the nucleotide T at each position can be read.

**Figure 3:**
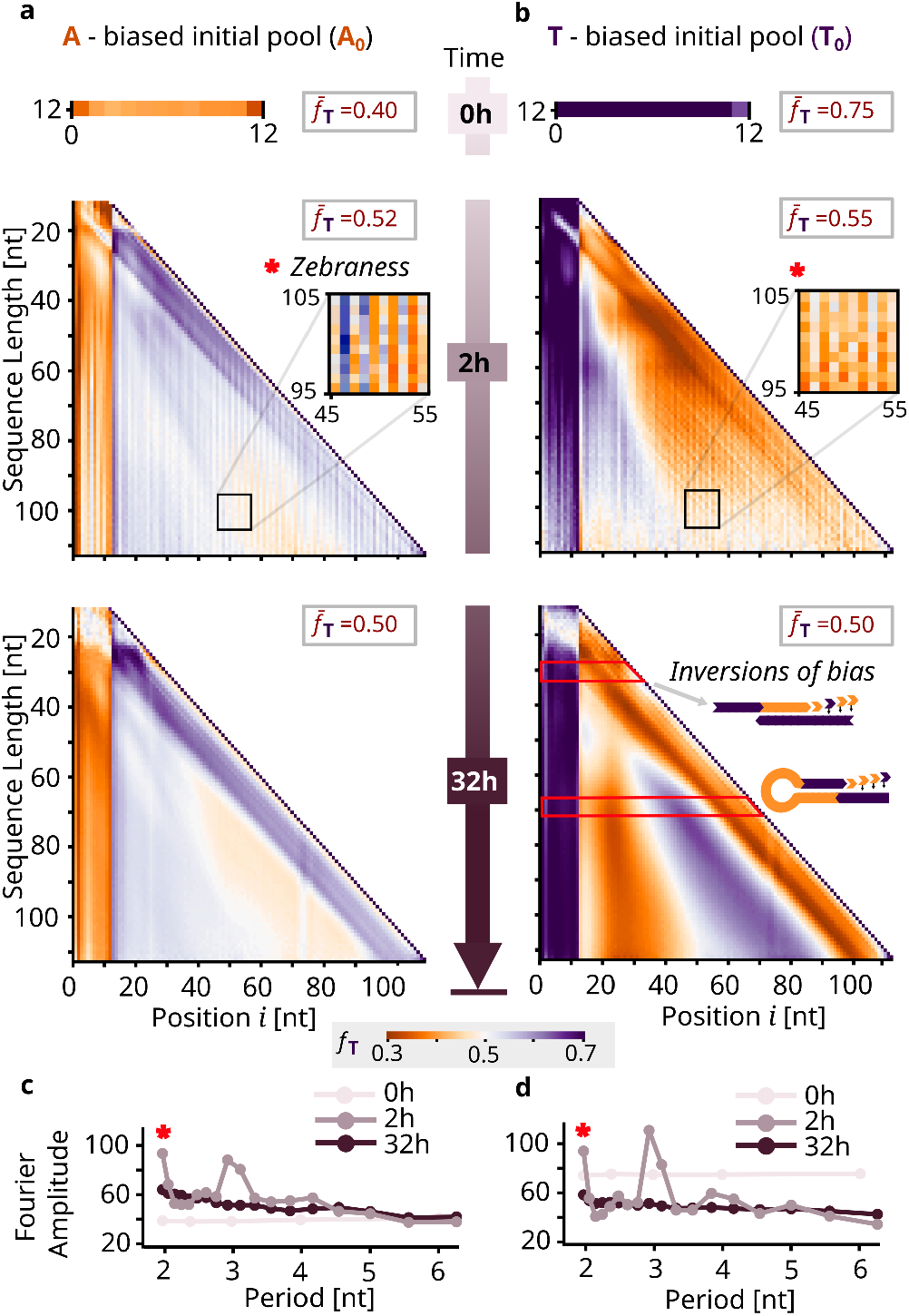
Effects of initial bias and elongation on sequence composition for AT (A_0_ and T_0_ experiments). **a, b** Evolution of nucleotide fraction *fT* across sequence lengths and positions in sequences for the initial pool, a middle time point and a final time point (0h, 2h, 32h). A-rich regions (*f_T_* ≪ 0.5) represented in orange and T-rich regions (*f_T_* ≫ 0.5) in purple. The initially biased average pool nucleotide fraction 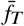 is countered as the pool undergoes polymerization, homogenizing to 0.5 at later time points. The first 12 nucleotides at the 5′ end retain the initial sequence bias for all graphs, due to the directionality of the polymerization mechanism (5′-3′). In addition, an inverse bias at the 3′ is explained explained by pool templation from the biased pool. For the 2h time point, horizontally alternating “zebra” patterns of *fT* are visible, illustrated by the insets with increased contrast. At 32h, gradients of alternating nucleotide fraction suggest self-complementarity, possibly a consequence of selftemplation, SI section XIII **c, d**. Periodicity is plotted as the amplitude of the Fourier modes of a discrete Fourier transform performed on the position-dependent conditional probability of A for the 50 mer long sequences, SI section XII C. The fast replicator sequences from the 2h middle time point display patterns with period 2 nt, matching the zebra patterns of the nucleotide fraction graphs, as well as period 3 nt.

The initial pools (Figure 3 **a** and **b**, top) consisted of 12 mer sequences with an overall T-fraction (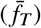 of 0.40 for A_0_ and 0.75 for T_0_. The heavier bias in the T_0_ pool can be explained due to DNA synthesis variability. Across positions, the distribution of the nucleotide fraction is close to homogeneous, with no apparent patterns in the initial pool that could propagate with replication.

The initial bias is countered by polymerization, and the overall pool average approaches equal nucleotide fraction for both A_0_ and T_0_ 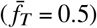. This can be seen for the 2h and 32h time points. As most of the sequences in the pool are biased towards one nucleotide, sequences are likely to find a similarly biased template. SI section IV shows the histograms of nucleotide fractions for all of the pools. Template-directed polymerization incorporates complementary nucleotides to the templates, inverting the bias in the newly forming strand segments. Note that all the sequences in the initial pool are 12 mer and that primer and template is a notation that solely depends on the direction of elongation, i.e. *Bst* adds nucleotides to the 3′ end of the primer, Figure 1.

While the pool-averaged bias was homogenized, in-strand positional biases were amplified. Due to the 5′-3′ direction of the polymerase, any bias at the priming first 12 nucleotides at the 5′ terminus will be preserved over the complete reaction period. Additionally, since the nucleotides added are mostly complementary, the nascent segment will be inversely biased. These bias inversions are observed as starting 12 mer columns for the 2h and 32h time points, for both of the analysed pools.

Fast replicators, corresponding to sequences observed at early time points, feature patterned structure. They display a zebra pattern, visible through the vertical stripes indicating alternating average nucleotide fractions, Figure 3 **a** and **b**, insets. Zebraness, defined as the fraction of alternating 2 mer motifs [38] (‘AT’ or ‘TA’ in this case), increases with sequence length for the 2h time points for both the A_0_ and T_0_ experiments, SI section X. It is thus a characteristic of fast replicators in AT experiments. An analysis of the 4 mer motif distribution supports similar conclusions: for fast replicators, alternating motifs such as ATAT and TATA are more abundant than bulky ones, which characterize the biased initial pools, SI section XI.

To understand the interdependency between in-strand sequence motifs, we calculated a matrix that correlates the nucleotide fraction at each position to all positions of each respective sequence for sequences of length 50, SI section XII.

The 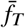 plots do not allow to do so as they average over all sequences of the same length. The correlation matrices for 2h time points, for both A_0_ and T_0_, revealed a diagonal correlation indicative of periodicity. To obtain the dominant period of the patterns, a discrete Fourier transform, SI section XII C, was applied to every row of the correlation matrices and averaged across all rows and sequences, Figure 3 **c** and **d**. The graphs spike at period 2 nt and 3 nt above the baseline Fourier amplitude of 50 which random sequences would display (the baseline equals the average pool nucleotide fraction in percent). As seen before, fast replicators favour high zebraness with a period of length 2 nt. Additionally, a periodicity of length 3 nt is revealed.

After 32h of polymerization, the zebra patterns in the fraction of T have been replaced by smooth gradients. A reason for this may be that the fast replicators have elongated even more and are no longer captured in the graphs. The gradients are antisymmetric around the center, corresponding to alternating inversions of bias. This indicates self-complementarity, suggesting self-templation through the formation of hairpins as a mechanism of elongation. Self-templation is favoured over pool templation when possible since it is kinetically more likely to find a complementary region within the proximity of the same molecule than within another molecule of the pool, Figure 1. Indeed the longest self-complementary region found in each sequence, averaged for each sequence length, is consistently longer in the replicated pools than in a randomly generated pool with *f*_*T, pool*_ = 0.50, SI section XIII. The 4 mer motif distributions support the same conclusion, as motifs that are reverse complementary have the same abundance, SI section XI. This selection for complementarity, through pool and self-templation mechanisms, leads to the stable convergence of the *f*_*T, pool*_ to 0.50, regardless of initial bias.

Similarly to the AT experiments, the G_0_ and C_0_ samples were analyzed with NGS, Figure 4. The three time points chosen in this case were adapted to the faster GC elongation kinetics. The initial pools had symmetric biases, with 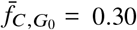 and 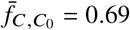 for C_0_. In the case of the polymerized pools, the sequences obtained were overall shorter than in the case of the AT data sets, even for the later time points. This may be due to a combination of the different polymerization dynamics and a lower sequencing efficiency for GC samples, SI section III, which yields fewer and lower quality reads for a similar initial concentration. For this reason, the GC graphs, Figure 4 **a** and **b**, are noisier and have shorter maximum length. For the earlier time point, at 0.5h incubation time, the alternating vertical stripes that indicated zebraness in the AT graphs are not present. In fact, when computing the zebraness for the GC samples, SI section X, the fast replicators, present at early time points, do not have significantly higher zebraness than sequences at the later time points. Overall, the zebraness of all GC samples is consistently below 0.5, indicating a majority of bulky 2 mer motifs (CC and GG). This result is confirmed by the 4 mer motif analysis, SI section XI.

**Figure 4:**
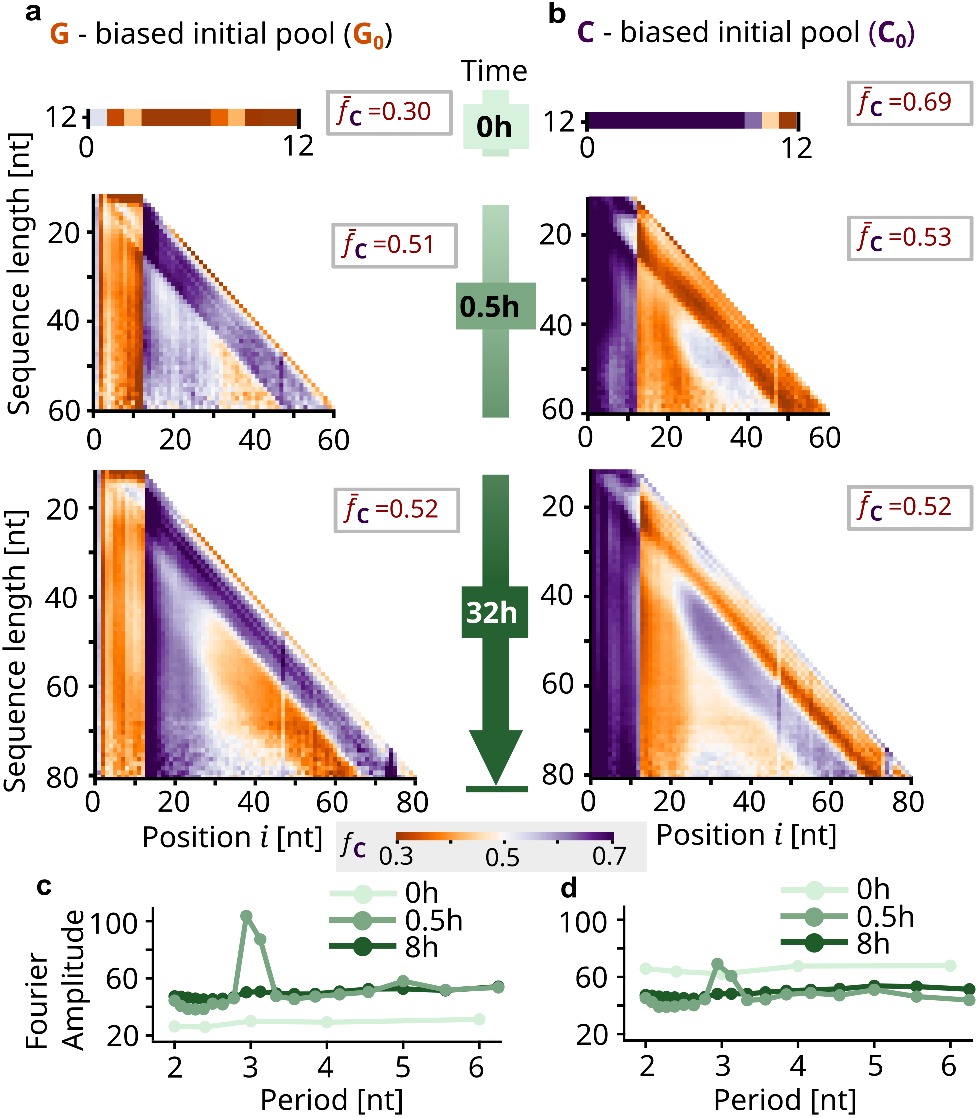
Results for GC (G_0_ and C_0_ experiments). **a, b** The nucleotide fraction *fC* for the 0h, 0.5h and 8h time points was decomposed by length and position, which leads to graphs similar to those for AT. Again, the initially biased average pool nucleotide fraction 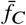 is homogenized with time and the first 12 nt retain the initial bias while the following segment is inversely biased due to pool templation. However, no zebra patterns are visible in the middle 0.5h time points. **c, d** The Fourier modes (for G, 50 mer) confirm this absence of 2 nt periodicity but do indicate 3 nt periodicity.

The difference between the AT and GC fast replicators can be explained by the intrinsic differences in the stacking energy of bulky and zebra motifs. Stacking energy of neighbouring nucleotide pairs is the main contributor for duplex stability [39]. For GC, zebra motifs are more stabilizing than bulky motifs [38], whereas for AT the opposite is true. The sequences rich in the most destabilizing motif type replicate the fastest into very long strands. This prevents them from being stuck in very stable secondary structures and renders them more accessible for several rounds of priming.

The inversion of bias both on the 5′ to 3′ end and in the intermediate region is evident for both the 0.5h and 8h time points as in AT. These can be explained with the pool and self-templation mechanisms. 4 mer motifs which are reverse complementary display similar abundances after polymerization has occurred, which can be seen by the symmetry in the graphs, SI section XI. The increase in overall pool complementarity leads to the convergence of the pool average nucleotide fraction to 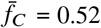 after eight hours for both experiments. Yet, this does not coincide with an increased length of the self-complementary regions in contrast to that of the AT pools, suggesting that long self-complementary stretches in the GC pools are not easily elongated and self-templation is not as favored, SI section XIII. The increased stability of the GC base pair with respect to the AT one may be the reason for strongly self-complementary sequences being stuck in overly stable duplexes, keeping them from further elongation. Coupling a strand-displacing system with environmental fluctuations could potentially retrieve these duplexes for further elongation.

The positional dependencies within sequences of a specific length were analyzed by conditional probability graphs which revealed periodicity in GC samples as they did for AT, SI section XII A. The Fourier transform graphs indicate an increased periodicity of 3 nt, Figure 4 **c, d**.

## Conclusion

We demonstrated that in following templated replication, pools display a positional bias and the average pool nucleotide fractions become more homogeneous. Replication from two independently synthesized initial pools with the same bias resulted in reproducible length distributions, average pool nucleotide fractions and sequence structure, SI section VI.

We experimentally verified that compositional diversity, represented by the average pool nucleotide fraction, arises from biased binary pools via templated replication. This is a necessary characteristic for the exploration of sequence space with the possibility of generating a functional sequence. Similar conclusions have previously been described for binary DNA systems *in silico* [10], particularly for templated ligation.

Simultaneously, the replication of an initially biased pool resulted in regions in the replicated sequence that possess the same or the symmetric bias, which alternate and balance each other on average. This allows for a biased exploration of subsections of sequence space with structured sequences, without restricting the sequence space to a subset of similar sequences. Different nucleotide biases have been shown to correlate with enrichment of different secondary structures [20], implying that the sequences obtained from our templated replication may exhibit a diverse range of secondary structure, which is in turn correlated with functionality.

Symmetry breaking, triggered by the selection for the reverse complement due to templation mechanisms, has been experimentally described for templated ligation. In a previous study [1] where binary AT pools were studied, two different sub-populations of sequences were found to contain a high amount of reverse complement sequences, with different nucleotide biases being enriched for each sub-population (an A-rich and a T-rich). Indeed, we observed a comparable behaviour within single sequences.

We also found that highly periodic sequences are replicated faster, interestingly amplifying a periodic trimer structure in all studied pools. Besides this agreement in 3 nt structure, the 2 nt periodicity differed for the two binary systems investigated. AT pools favored 2 nt zebra motifs such as AT and TA, whereas GC pools preferred bulky motifs such as GG and CC. Our findings, especially of the high self-complementarity in long AT sequences, SI section XIII, support the mechanism of “hairpin elongation” for repetitive DNA, as previously suggested [40]. Repetitive DNA strands possess a high number of potential fold-back sites for hairpin formation. Repeated complete or partial melting, possibly induced by the strand displacing activity of *Bst*, alternating with hairpin formation and self-templation, would quickly elongate highly repetitive sequences.

In this study, we employed an experimental model system to provide insight into the role of replication as a mechanism of selection. Using a protein-based replication system with strand displacement (*Bst*), we identified which sequence patterns emerged as the fittest by analysing the fast replicators. In addition, we characterized the dependency of the emergent structure on the initial pool. Overall, our findings contribute to elucidate the steps involved in the Darwinian evolution of short unstructured nucleic acids into long functional sequences.

## Methods

### Polymerization with *Bst*

The polymerization reactions were performed with *Bst* 2.0 DNA Polymerase (New England BioLabs). The conditions were according to the protocol provided by the manufacturer: 1x Isothermal Amplification Buffer, 8mm MgSO_4_ (for a total of 10mm with the 2mm MgSO_4_ from the 1x buffer), 320U/mL *Bst* (all supplied when ordering the enzyme), 1.4mm of each nucleotide triphosphate and 10μm DNA. AT samples were supplied with 1.4mm dATP and dTTP and GC experiments with 1.4mm dGTP and dCTP (all from Sigma-Aldrich). All experiments were conducted with initial DNA samples containing only random 12 mers provided by biomers.net, with binary base alphabets (AT, GC) in varying base content, SI table I.1. The ordered base content differs from the effective base content detected with NGS, Figure 3, 4 and SI section VI. The polymerization reactions were incubated in a thermocycler with the following protocol: 1. constant temperature (35°C for AT, 65°C for GC) for the reported time; 2. 90°C for 20min to deactivate *Bst*. The incubation temperature was lower for AT than for GC due to differences in melting temperature, and based on a temperature screening performed with *Bst*, SI section VII.

### PAGE and gel imaging

The samples were run in a denaturing 15% polyacrylamide in 50% urea, with a 19:1 ratio of acrylamide to bis-acrylamide and polymerized with TEMED (tetramethylethylenediamine) and ammonium persulphate. The gels were pre-heated in the electrophoretic chamber at 300V for 27min. The samples were then loaded, in a mixture with a ratio of 2:7 of sample to loading dye. Loading dye is prepared in-house (for 10mL: 9.5mL formamide, 0.5mL glycerol, 1μL EDTA (0.5m) and 100μL Orange G dye (New England BioLabs). The samples were at 50V for 5min followed by 300V for 25min. After the run, the gels were stained with a 2x SYBR Gold (Thermo Fischer Scientific) dilution in Tris-Borate-EDTA buffer 1x. They were then rinsed with 1x TBE buffer twice and imaged using a BioRad ChemiDoc™ MP imaging system. The 20-100bp ladder (DNA oligo length standard 20/100 Ladder, IDT) was supplied in a final concentration of 2.04ng/μL (for each rung) and the 100-1000bp ladder (100bp DNA Ladder, New England BioLabs) in a final concentration of 71.4ng/μL (for all rungs; concentrations vary by *n* mer as described by the manufacturer). Finally, the obtained micrographs were loaded into and analyzed with a self-written LabVIEW program, SI section IX.

### Sequencing

Samples were sequenced by the Gene Center Munich (LMU) using the NGS Illumina NextSeq 1000 machine (flow cell type P2, 2 × 50bp with 138 cycles for 100bp single-end reads; at most 120bp with 2 indexes were read, with declining quality towards the end). 50 million reads were ordered for each sample. The raw sequencing data obtained, in FastQ format, was processed in this order by demultiplexing, quality score trimming, and regular expression filtering. Demultiplexing was performed with software from Galaxy servers [41], provided by the Gene Center Munich. During sequencing, each read base was assigned a Phred quality score *Q* = −10 log_10_ *P*, where *P* is the probability of an incorrectly read base [42]. Using Trimmomatic [43] we trimmed low quality segments by running a sliding window of 4 nt in the 3′ to 5′ end direction over the sequence that allowed a minimum average Phred quality of 20, otherwise trimming at the leftmost base of the window, corresponding to an average accuracy of at least 99%. As the experimentally obtained sequences were appended on the 3′ terminus with a CT-region followed by an AGAT during sequencing preparation, those needed to be found and cut, for which we employed the following regular expressions:

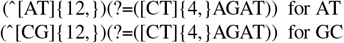

This also ensured that only binary sequences were included in the analysis.

## Data availability

All data and code relevant to the study are included in the paper, uploaded as supplementary information or, in the case of the raw sequencing FASTQ data, provided in the external data repository xxx.

## Acknowledgment

The authors acknowledge support from the European Research Council (ERC Evotrap, Grant Number 787356), the CRC 235 Emergence of Life (Project-ID 364653263), the Excellence Cluster ORIGINS which is funded by the Deutsche Forschungsgemeinschaft (DFG, German Research Foundation) under Germany’s Excellence Strategy – EXC-2094– 390783311, and the Center for NanoScience (CeNS). We also thank Christof B. Mast, Sreekar Wunnava and Paula Aikkila for comments on the manuscript, Annalena Salditt for helpful discussions on data analysis, as well as Stefan Krebs and Marlis Fischalek at the Gene Center Munich for their help with the library preparation and the sequencing of the samples.

## Author contributions

Project conception: A.C.S., D.B., Research design: A.C.S., F.T.D., D.B., Methodology development: A.C.S., F.T.D., Experiments: A.C.S., F.T.D., Z.M., Data analysis: A.C.S., F.T.D., Programming: F.T.D., Manuscript writing: A.C.S., F.T.D., Manuscript reviewing: A.C.S., F.T.D., D.B., Supervision: A.C.S., D.B. Funding acquisition: D.B.

## Additional information

The online version contains supplemen tary material available at https://…

