## Supplementary information for "Replication elongates short DNA, reduces sequence bias, and develops trimer structure"

Adriana Calaça Serrão,[1,*] Felix T. Dänekamp,[1,*] Zsófia Meggyesi,[1] and Dieter Braun[1,†]

[1]*Systems Biophysics, Physics department, Center for NanoScience, Ludwig-Maximilians-Universität München, Amalienstraße 54, 80799 Munich, Germany*

#### Contents

I. DNA Strands 2

II. Bst enzyme 2

III. Sequencing yield and read length distribution 2

IV. Nucleotide fraction distribution 4

V. PAGE images with elongation kinetics for all pools 4

VI. Reproducibility 5

VII. Choice of experiment conditions 5

VIII. Smear quantification in gels 6
- A. PAGE and smears 6
- B. A model for smear quantification 6
- C. Small second order changes of intensities by length 7
- D. Required measurements: intensities and ladder peak positions 7
- E. Total molar concentrations 7

IX. Extraction of concentrations and length distributions from PAGE images 8

X. Zebraness 8

XI. Motif analysis 8

XII. Probability graphs and sequence structure 10
- A. Conditional probabilities 10
- B. Average probabilities and length evolution 10
- C. Fourier analysis and periodicity 10

XIII. Self-complementarity 10

---

[*] equal contribution

[†]

### Supplementary information I: DNA Strands

| Pool | Sequence |
|---|---|
| G-biased | 5′ SSSSSSSSSSSS(67%G:33%C) 3′ |
| C-biased | 5′ SSSSSSSSSSSS(33%G:67%C) 3′ |
| A-biased | 5′ WWWWWWWWWWWW(67%A:33%T) 3′ |
| T-biased | 5′ WWWWWWWWWWWW(33%A:67%T) 3′ |
| A-biased* | 5′ WWWWWWWWWWWW(50%A:50%T) 3′ |

**Table I.1:** DNA sequences as ordered from biomers.net. S signifies GC and W AT. The actual nucleotide content of the obtained initial pools is shown in section IV.

### Supplementary information II: Bst enzyme

*Bacillus stereathermophilus* polymerase I (Bst) is a thermophilic, strand displacing A family polymerase used in many isothermal amplification applications [34]. The enzyme binds to a double stranded segment and elongates it in the 5′-3′ direction, adding the nucleotides complementary to the template and displacing any other downstream bound strands.

Specifically, in this work, Bst 2.0 DNA Polymerase was used. This is an *in silico* designed homologue of the large fragment of the *Bst* enzyme that contains the 5′-3′ polymerase activity, but lacks 5′-3′ exonuclease activity. It also has higher speed, yield, salt tolerance and thermostability than the wild-type. This enzyme should have an activity of about 10% for 35°C and 100% for 65°C, and has maximum performance at 4-10mM $Mg^{2+}$ according to the manufacturer (New England Biolabs).

The large fragment of *Bst* has proofreading activity which contributes to its high fidelity. This is not achieved through an exonuclease domain, but though a mechanism that checks the structure of the incorporated nucleotide at the active site [32]. The active site of the protein interacts with the minor groove of the double stranded DNA helix, particularly the 4 base pairs closest to the 3′-OH terminus. The proofreading is done at the last 3′ base. In case the base added is wrong (i.e. not complementary to the template), the 3′-OH terminus will not be oriented correctly and the elongation will not proceed [44].

In addition to the downstream bound strands, any secondary structure of the primers or template is denatured by *Bst* due to its strand displacing activity – it does not ‘slip’ [45].

### Supplementary information III: Sequencing yield and read length distribution

The length distribution of the reads obtained after quality processing, adapter trimming and regular expression filtering, as described in the methods section, is plotted in Figure III.1. For all of the samples, the initial pool is almost entirely composed of 12 mer strands. Shortly after the onset of elongation, in the middle time points (2h for AT data sets and 0.5h for GC data sets), the amount of 12 mer strands decreases, as they have been recruited for replication. The middle time point curves peak at lengths between 20 and 30 nt, similar to the length distributions obtained via PAGE smear quantification, Figure 2. The end time points reveal a depletion both of the initial 12 mer and of the 20-30 nt peak, indicating that both 12 mer and longer strands get recruited in the later stages of replication. This supports the idea that one strand can go through several rounds of replication.

The maximum read length obtained with Illumina NGS was 112 nt (after cutting the CT-tail and AGAT), which is the reason for the sharp wall in the AT datasets. There are likely longer products only partially sequenced, which are discarded by regular expression filtering. The longer strands are visible in the PAGE images for the AT data sets, Figure 2 and section V. The most striking difference between the length distributions obtained for the two methods (PAGE and NGS) is the higher abundance of long (up to 112 nt) products compared to short ($\approx$ 12 mer) in the case of NGS. This favoring of longer strands seems to be a systematic bias of sequencing. Since we know that the amount of strands should stay the same (10µM total DNA strands), the read counts should be constant across all data sets, which is not the case, Figure III.2. Indeed, the counts observed were about two orders of magnitude lower for the initial time points. This could be due to a reduced yield for the adaptor ligation step depending on the fragment length.

Additionally, GC data sets have less counts than AT data sets. As this is already the case for the initial 12 mer pools, likely is due to differences in sequencing yield possibly related to challenges in amplification. For instance, both very high and low GC contents (both the case in the work as the systems were binary) have been shown to be a challenge to NGS due to difficulties in the PCR amplification step [46].

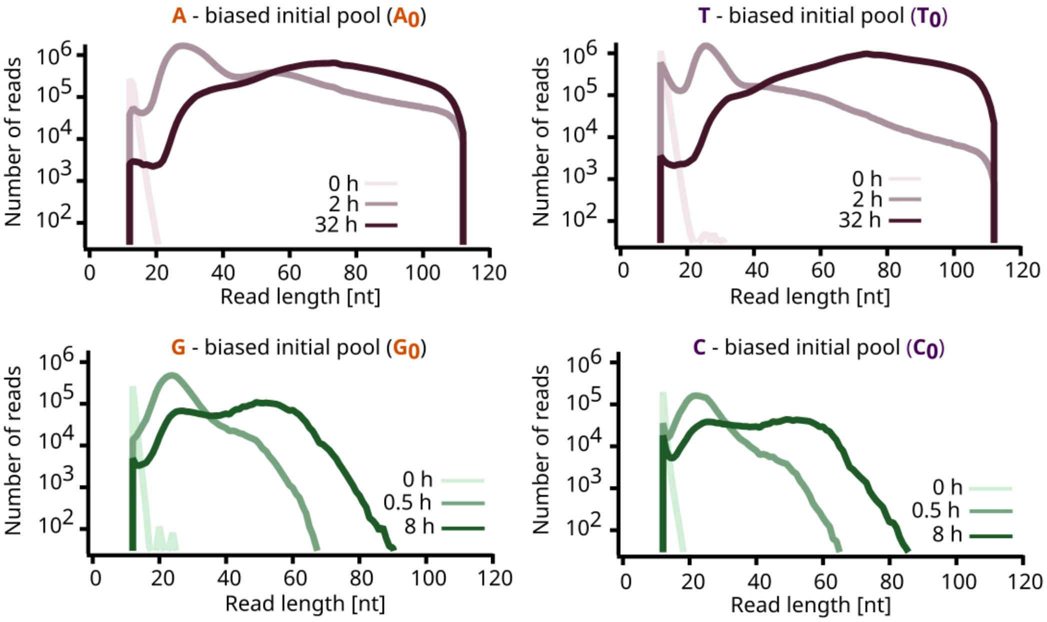


**Figure III.1:** Length distribution of the NGS reads obtained for each of the data sets, after pre-processing. The initial pools are composed almost entirely of 12 mer strands. Those strands are depleted as sequences get recruited for replication. Between the middle and the late time point, 12 mer and longer strands are depleted, supporting the idea that one strand can go through several rounds of replication. Strands longer than the maximum read length of 112 nt are only partially sequenced and were discarded during pre-processing.

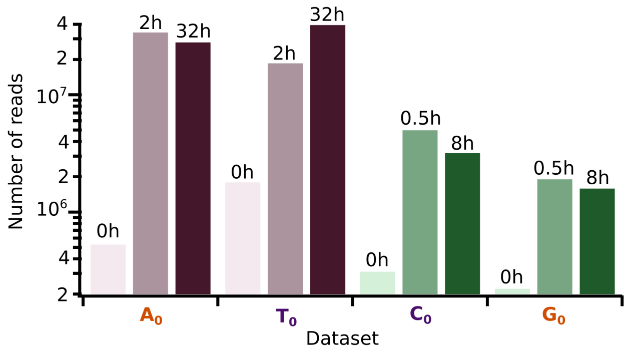


**Figure III.2:** Total NGS read counts obtained for all the data sets, after pre-processing. As the amount of strands (10µM total DNA) should in theory remain the same, sequencing is revealed to be systematically biased, favoring longer strands. Additionally, GC data sets systematically have less read counts than AT ones, which likely is due to differences in sequencing yield.

### Supplementary information IV: Nucleotide fraction distribution

The nucleotide fraction of all sequences in each of the data sets was computed and plotted in Figure IV.1. The initially biased pools shift towards a distribution centered around 0.5, corresponding to a homogeneous average pool nucleotide fraction. There is no significant difference between the middle time point and the later time point, indicating a rapid reduction of bias.

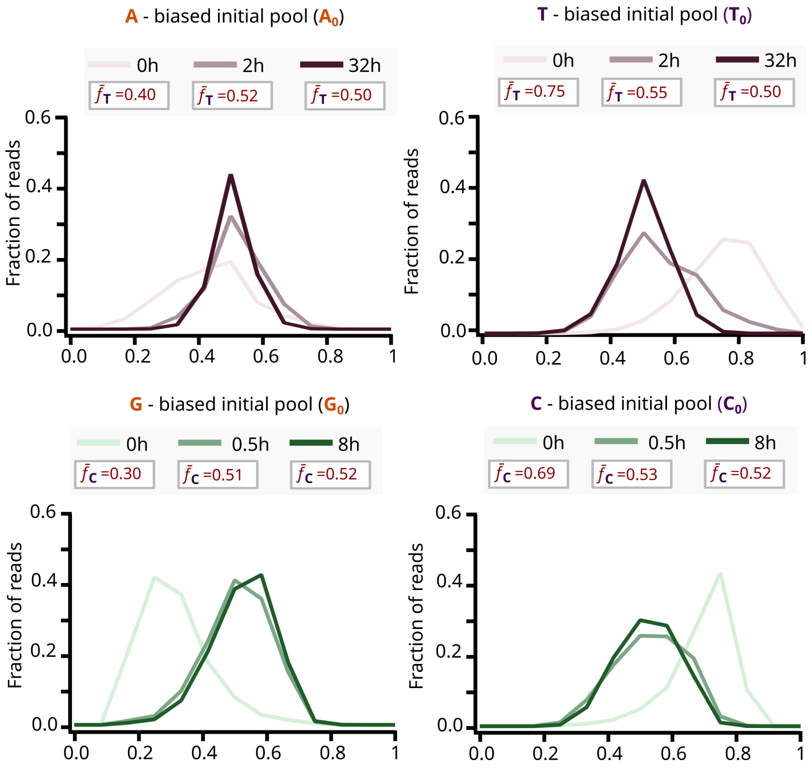


**Figure IV.1:** Distribution of nucleotide fraction among the obtained reads after pre-processing for all the sequenced pools and time points. The initially biased pools shift towards a distribution centered around 0.5, corresponding to a homogeneous average pool nucleotide fraction. Middle and late time point distributions are very similar, indicating a rapid reduction of bias.

### Supplementary information V: PAGE images with elongation kinetics for all pools

Figure V.1 displays the PAGE images for all replication experiments including the inversely biased pools not previously presented in Figure 2. The A*$_0$ experiment independently reproduces the A$_0$ one, see section VI.

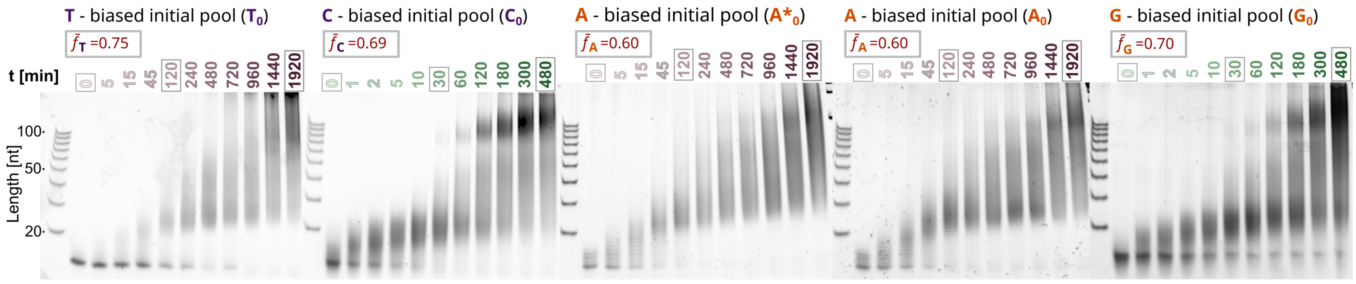


**Figure V.1:** PAGE images for all replication experiments from the main paper, Figures 3 and 4, as well as the reproducibility experiment, section VI.

### Supplementary information VI: Reproducibility

Reproducibility of the obtained length distributions from polymerization was checked with different aliquots of the initial pools in similar experiments, leading to similar PAGE images. While reproducibility was not purposefully investigated by sequencing these aliquots, we did obtain two initial pools similar in average base nucleotide fraction when trying to perform experiments with a non-biased AT pool (50%A, 50%T). After sequencing, the base content was revealed to be similar to the $A_0$ pool, providing us with an independent repeat. The PAGE image for this data set labeled as $A^*_0$ is shown in section V alongside $A_0$, displaying a very similar evolution of the length distributions. The graphs depicting average probabilities to find a base at a certain position are given in Figure VI.1, where the graphs for $A_0$ from the main paper are reproduced alongside it. The prominent resemblance is a good indicator of general reproducibility not only for length distributions and average pool nucleotide fractions, but also for the structure of sequences. The Fourier transform reveals the same periodicities of 2 and 3 nt.

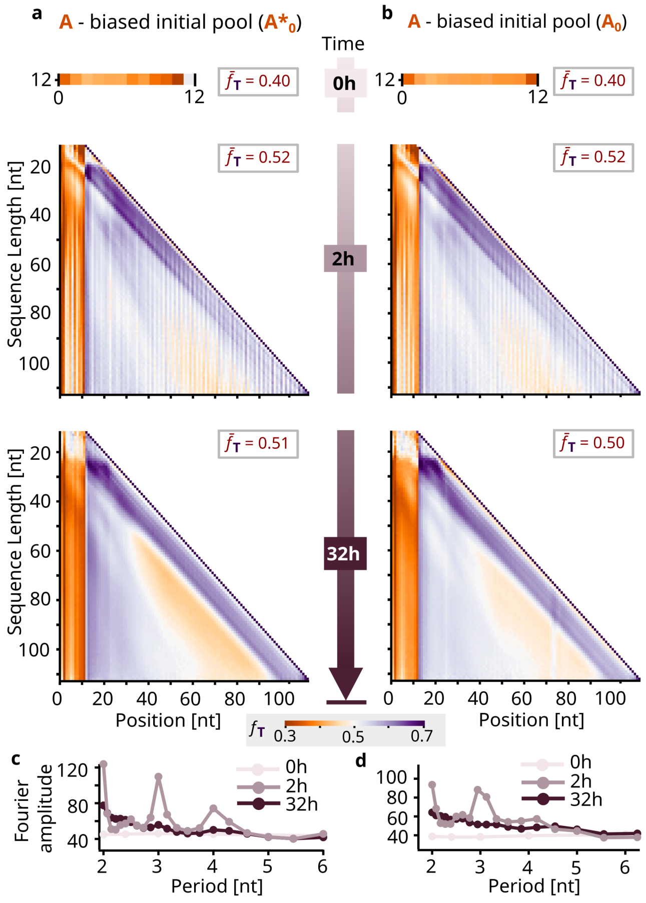


**Figure VI.1:** Reproducibility on the sequence level: High similarity between the independent $A^*_0$ and $A_0$ graphs for all timepoints. From similarly biased initial pools, very similar elongated pool structure emerges. The Fourier transform reveals the same periodicities of 2 and 3 nt. The PAGE image for the reproducibility experiment is given in section V, displaying a very similar evolution of length distributions.

### Supplementary information VII: Choice of experiment conditions

Previous to the time point experiments we performed a temperature screening varying the temperature between 35°C-45°C for the AT pools, and 45°C-75°C for the GC pools. The temperatures of 35°C (AT) and 65°C (GC) for the final experiments were chosen because the screening indicated these were the optimal temperatures for the *Bst*, i.e. the products obtained were longer. PAGE images from temperature experiments are provided in Figure VII.1. The lower efficiency of polymerization in these experiments compared to the final time point experiments presented in section V is likely due to a different dNTP content at this stage of screening (14 mM here versus 1.4 mM).

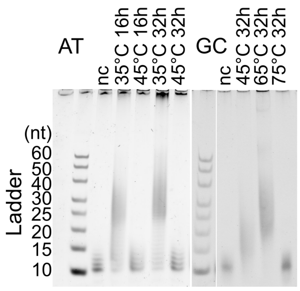


**Figure VII.1:** PAGE images for prior experiments with varying temperatures. 35°C for AT and 65°C for GC were chosen as optimal temperatures for fast polymerization by efficient strand displacement. The lower efficiency of polymerization in these experiments compared to the final time point experiments presented in section V is likely due to a different dNTP content at this stage of the screening (14 mᴍ here versus 1.4 mᴍ).

### Supplementary information VIII: Smear quantification in gels

#### A. PAGE and smears

Quantification of PAGE images, that is extracting the concentration of DNA of certain lengths from the gel image, is difficult for two reasons. Firstly, the functional relation of imaged intensity of fluorescence to the concentration of DNA in the gel is usually unknown, though for SYBR gold, fluorescence signal was found to increase linearly with stained DNA concentration as well as DNA strand length for a wide range of concentrations [47]. Secondly, DNA strands of the same length move through the gel with a slightly different speed due to diffusion and influence of sequence composition, leading to a smearing of signals through an overlap of n mer bands (DNA strands with a length of $n$ nt) with neighboring $n \pm \Delta n$ bands. As polymerization, unlike for example ligation, produces DNA strands of all possible discrete lengths which result in continuous intensity smears in gels, a direct summation of the intensity signal for DNA strands of a specific length is impossible.

#### B. A model for smear quantification

Instead, a model relying on small second order changes in intensities of neighboring DNA strands is developed. The central idea of the model is that for obtaining the area $A$ of a centered symmetric peak, e.g. a Gaussian peak, instead of integrating the peak function from $-\infty$ to $\infty$, it is possible to integrate the sum of infinite evenly spaced equally shaped peaks from $x - \frac{1}{2}\Delta\mu$ to $x + \frac{1}{2}\Delta\mu$ with peak centers spaced with $\Delta\mu$ (1):

$$A = \int_{\mathbb{R}} G_m(h_m, \sigma_m, \mu_m)\,\mathrm{d}x = \int_{\mu_m - \frac{1}{2}\Delta\mu}^{\mu_m + \frac{1}{2}\Delta\mu} \sum_{n=-\infty}^{\infty} G_n(h_m, \sigma_m, \mu_n)\,\mathrm{d}x \tag{1}$$

where $h$ describes the height, $\sigma$ the width and $\mu$ the position of the peak $G(h, \sigma, \mu)$.

As peaks are centered, one can then neglect contributions from the sides with little error and consider only the immediate neighboring peaks (2):

$$A \approx \int_{-2\sigma_m}^{2\sigma_m} G_m(h_m, \sigma_m, \mu_m)\,\mathrm{d}x \approx \int_{\mu_m - \frac{1}{2}\Delta\mu}^{\mu_m + \frac{1}{2}\Delta\mu} \sum_{n=-\Delta m}^{\Delta m} G_n(h_m, \sigma_m, \mu_n)\,\mathrm{d}x \tag{2}$$

with $\Delta m$ such that $|\mu_{m\pm\Delta m} - \mu_m| \approx 2\sigma_m$.

The inner part of the integral resembles the measured gel intensity $I(x)$ at point $x$, which is described by (3):

$$I(x) = \sum_{n=-\Delta m}^{\Delta m} G_n(h_n, \sigma_n, \mu_n) \tag{3}$$

when gel bands are modeled as centered symmetric peaks. The only difference is that in the gel, neighboring peaks are not evenly spaced and are described by different $h$'s and $\sigma$'s (observe that $h_m$ and $\sigma_m$ were replaced by $h_n$ and $\sigma_n$).

##### C. Small second order changes of intensities by length

But how large is the error, if neighboring gel peaks are assumed to be evenly spaced and equally shaped? – Very little error can be associated with the variability of peak spacing in the immediate vicinity of each peak. For the peak heights and widths, this doesn't hold. Deviations would lead to larger errors.

In most cases though, while intensities of neighboring gel bands vary, the change in intensities doesn't vary much. But when second order changes are small, peaks of same distance to the one that is to be measured can be described with symmetric deviations in $h$ and $\sigma$ (4) and (5):

$$G_{m-\Delta m} = G(h_m \pm \Delta h_m, \sigma_m \pm \Delta\sigma_m, \mu_{m-\Delta m}) \quad (4)$$

$$G_{m+\Delta m} = G(h_m \mp \Delta h_m, \sigma_m \mp \Delta\sigma_m, \mu_{m+\Delta m}) \quad (5)$$

In this case, the relative error in the area calculation (2) can expected to be smaller than $\frac{2\Delta h_m \Delta\sigma_m}{h_m \sigma_m}$ and should stay reasonably small unless intensities change abruptly with certain oligomer lengths.

Assuming a continuous smear with slow second order change in intensity, total intensities $A_m$ of m mer peaks are approximated well by (6):

$$A_m = \int_{\mu_m - \frac{1}{2}\Delta\mu}^{\mu_m + \frac{1}{2}\Delta\mu} I(x)\,\mathrm{d}x \quad (6)$$

##### D. Required measurements: intensities and ladder peak positions

The required values remaining besides the gel intensity $I(x)$ are then the position of the m-th peak, $\mu_m$, and its spacing to neighboring peaks, $\Delta\mu$, which was assumed to be constant in its vicinity and can thus be calculated as (7):

$$\Delta\mu(m) = \frac{\mu_{m+1} - \mu_{m-1}}{2} = \frac{\Delta\mu_{\pm m}}{\Delta m} \approx \frac{\partial\mu(m)}{\partial m} \quad (7)$$

with the continuous interpolation $\mu(m)$ of $\mu_m$'s, leaving only the function $\mu(m)$ relating oligomer lengths to gel positions to be found.

To determine $\mu(m)$, DNA ladders can be used: by interpolating the ladder rung mer peak positions of known oligomer lengths with a function relating in-gel-distances to product lengths, $\mu(m)$ is acquired. The simplest function yielding a good fit is a logarithm scaled by $m$, $\mu(m) = a\frac{\ln(bm)}{m}$, with fit parameters $a$ and $b$. For fitting of long strands (>100nt), another logarithmic factor improved the fit even more, giving the final fit function (8):

$$\mu(m) = a\frac{\ln(m)}{m} + b\ln(m) + c \quad (8)$$

with peak position in gel $\mu$, oligomer length $m$ and fit parameters $a$, $b$ and $c$. Fits were performed using the Levenberg-Marquardt algorithm implemented in LabVIEW.

##### E. Total molar concentrations

The accuracy of the PAGE gel quantification, should the total molar concentrations of DNA strands in each lane be known, can finally be checked by comparing calculated total molar concentrations to the known total molar concentration of the probes. The molecular weight of a single-stranded polymerized DNA oligomer can approximately be calculated as (9):

$$\text{molecular weight } [g/mol] = n * 308.95 - 61 \quad (9)$$

where $n$ is the number of nucleotides in the n mer. 308.95 is the average molecular weight of an incorporated nucleotide (A: 313.2, T: 304.2, C: 289.2, G: 329.2) and $-61$ is obtained by subtracting one phosphate (weight 79) for the hydroxyl-end and adding one water molecule (weight 18).

Reversing this line of thought, intensities can also be related to concentrations for each lane individually, as long as their molar concentrations are known. This normalization for each lane additionally helps to account for experimental variations in intensities introduced through pipetting of low volume / high viscosity samples and possible evaporation effects in thermocyclers. For these reasons, each gel lane was indeed normalized to have the total molar concentration (the sum of molar concentrations for all oligomer lengths) equal the known molar concentration of the initial sample for gel analysis in this work – the total molar concentration was known to equal the initial molar concentration at each time because elongation of initial DNA strands doesn't change total molar concentration.

### Supplementary information IX: Extraction of concentrations and length distributions from PAGE images

PAGE images were analyzed with a self-written (adapted from an existing program of the AG Braun) LabVIEW tool, which allowed to obtain the concentrations of DNA strands depending on length from smears in the gel lanes by using known total molar concentrations of each lane and the linear increase in fluorescence intensity of SYBR gold with strand length and concentration [47]. The total molar concentration is known since it stays constant throughout the experiment as no new strands may appear through polymerization. Effects of hydrolysis should be small. The analysis happened in three main steps: gel image to lane data conversion, ladder peak detection and concentration analysis.

Firstly, the PAGE image was converted to lane data by loading the image and extracting the intensities, matching a lane mask with the imaged lanes and performing a background correction. A uniform x-axis for all lanes was then calculated and the y-axis rescaled to make them comparable.

In the second step, the approximate positions of the ladder n mers in the leftmost lane of the gel (ladder peak positions) were detected. The program then fitted each peak with a Gaussian function individually to obtain the precise ladder peak position and interpolate the positions to arrive at a position vs. length function $\mu(m)$ as explained in section VIII.

In order to quantify the concentrations in the third step, an intensity to concentration relation was obtained for each gel lane individually by knowledge of the constant total molar concentration. The width function $\Delta\mu(m)$ was fixed to be the derivative of the position function as derived in subsection VIII D. This had the advantage of precisely covering the whole range when calculating concentrations for all n mers and permitted to obtain a measure of total molar concentration for each lane allowing a normalization of each lane for correction of experimental variability and better comparability.

### Supplementary information X: Zebraness

Zebraness has been found to be a useful measure of pool structure in a theoretical study simulating elongation of random RNA strands, permitting an investigation of evolutionary dynamics in sequence space [24]. Zebraness $\zeta(S)$ of a strand $S$ of length $L_S$ is defined as the number of alternating "zebra" motifs (XY or YX) within the sequence divided by the total number of binary motifs, given by $L_S - 1$. A random sequence is expected to have a zebraness of 0.5, a fully alternating strand one of 1 and a homogenous strand one of 0.

We define a system zebraness $Z_L$ of strands $S_L$ of the same length $L$ as the average of all $\zeta(S_L)$. Plotting $L$ vs. $Z_L$ reveals the lengths at which zebraness starts to become eminent in the pool, Figure X.1. There is an observed trend of enhanced zebraness in AT and suppressed zebraness in GC pools, a pattern that is reinforced as sequences get longer through replication.

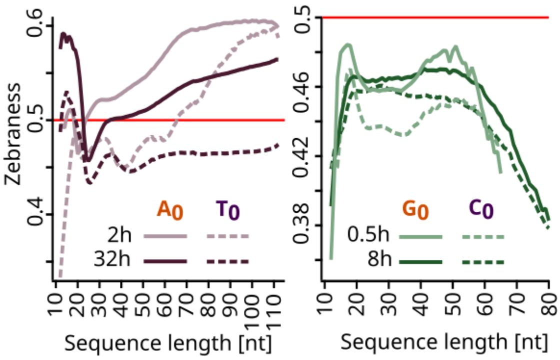


**Figure X.1:** Zebraness $Z_L$ by length, corresponding to fraction of 2 mer motifs that are XY or YX. In the case of AT samples (left panel) the intermediate time point samples (2h) have a higher average zebraness than their later time point counterparts. Additionally, the zebraness is higher for longer sequences, reinforcing that the 2 mer periodicity is present in fast replicators. In constrast, GC samples (right panel) have a generally lower zebraness, consistently below 0.5. Furthermore, the zebraness decreases for longer strands, indicating that the bulky 2 mer motifs are favoured for the fact elongators in the GC samples.

### Supplementary information XI: Motif analysis

A natural way to investigate sequence structure is to count the occurrences of n mer motifs in the whole sequence pool or parts of it. For each sequence, it is counted how often the n mer motif occurs inside of it with a sliding window starting at positions

0, 1, 2, ... Results are then summed over the selected pool part, yielding average trends of that pool part. Randomly generated sequences would yield almost perfectly evenly distributed motifs, as deviations quickly disappear through statistical narrowing.

A 4-motif analysis has high chances to reveal short-scale structure, which complements measures of zebraness as well as the probability analysis discussed in section XII. Recalling the different elongation mechanisms at play for shorter and longer strands as well as the differentiation of the pool in "fast replicators" and "stalled" sequences, the analysis is conducted for specific parts of the pools. In the 2h timepoint for AT and the 0.5h timepoint for GC, only sequences with a length of at least 40nt are analyzed whereas the 32h (AT) and 8h (GC) pool is split into two subsets, of which the 12-40nt and the 70-120nt (AT) or 40-120nt (GC) are displayed in Figures XI.1 and XI.2.

The symmetry in the 4 mer motif graphs reveals the enrichment in reverse complementary motifs for both pools after replication. The main difference between AT and GC is that the favoured motifs are more zebra-like for AT and more bulky for GC.

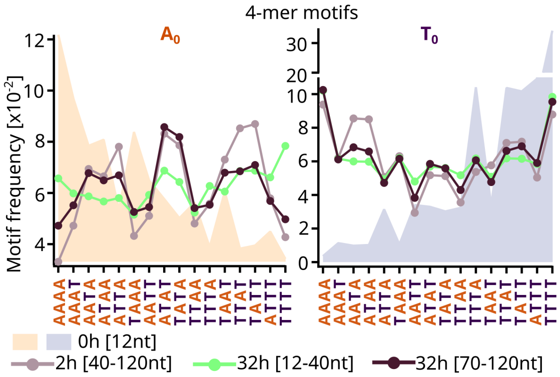


**Figure XI.1:** 4-motif distributions for $A_0$ (left panel) and $T_0$ (right panel) pools, at different time points. The motif distribution of the initial pool, in a solid color in the background, reveals a highly skewed distribution, due to the initial bias of the pool. For the 2h time point, the motifs were plotted for the sequences that are above 40 nt in length in order to characterize the fast replicators. These showed an enrichment in alternating motifs. For the 32h pool, two different length frames were analysed: 12-40nt and 70-120nt. The longer sequences had a similar motif distribution to that of the fast replicators, whereas the shorter sequences had a flatter distribution. The symmetry of the distribution indicates that reverse complementary motifs are enriched similarly.

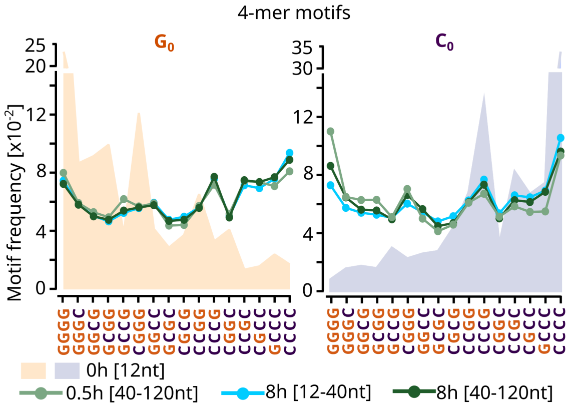


**Figure XI.2:** 4-motif distributions for $G_0$ (left panel) and $C_0$ (right panel), at different time points. The motif distribution of the initial pool, in a solid color in the background, reveals a highly skewed distribution, due to the initial bias of the pool. For the 0.5h time point, the motifs were plotted for the sequences that are above 40 nt in length in order to characterize the fast replicators. For the 32h pool, two different length frames were analysed: 12-40nt and 40-120nt. All of the replicated pools, regardless of the elongation time and length frame, had a rather homogeneous distribution of motifs, though there seems to be a slight preference towards bulky motifs.

### Supplementary information XII: Probability graphs and sequence structure

##### A. Conditional probabilities

To investigate sequence structure, conditional probabilities have proven useful. Selecting sequences of one length from the pool, we plot the probability of finding one of the two possible bases at a specific position depending on the base found at another position in a square graph, Figure XII.1.

The diagonal always displaying a probability of 100% (as that base is fixed by the analysis), two main methods of observing structure propose themselves. Vertical homogeneities correspond to structure present in the whole pool, as apparent in the first twelve positions of base $ii$, where the initial bias is clearly visible. Diagonal lines parallel to the main diagonal correspond to structure present in sequences individually, where the periodicity is shared across the pool but the onset of periodic patterns is not.

##### B. Average probabilities and length evolution

Whole pool structure from vertical homogeneities is best observed by averaging the conditional probabilities from the previous section vertically, that is with fixed position $ii$, obtaining a single horizontal line. This collapsing permits to plot averaged conditional probabilities for every length present in the pool, yielding a triangular graph illustrating gradients that point to an evolution of sequence structure with length and time. These graphs were given in the main paper for the three sequenced timepoints (including the zero one) for all four data sets.

##### C. Fourier analysis and periodicity

The presence of sequence periodicity, with onsets or shifts varying across the pool, visible as diagonal lines parallel to the main diagonal in conditional probability plots, suggests Fourier analysis as a method of quantification. For discrete one-dimensional sets of data, a discrete Fourier transform is defined as (10):

$$y_k = \sum_{n=0}^{N-1} x_n e^{\frac{-i2\pi}{N} kn} \tag{10}$$

where $x$ is the input data with $N$ elements and $y$ the transformed result. The amplitude $A_k$ of the sinusoidal component $e^{\frac{-i2\pi}{N} kn}$ of $x_n$ with the frequency $f_k = \frac{k}{N}$ encoded by the complex $y_k$ is given by $A_k = \frac{|y_k|}{N}$. The period $p_k$ is the inverse of the frequency $f_k$, $p_k = \frac{1}{f_k} = \frac{N}{k}$. Fourier transforms in our analysis were performed with the fast Fourier transform algorithm provided by LabVIEW.

From the conditional probability data discussed in subsection XII A, periodicity was investigated by separately performing a discrete Fourier transform for each horizontal lane of data with fixed position $i$ and then summing the obtained amplitudes $A_k$. As the Fourier transform is linear and the phase shift is dismissed, this recovers the frequencies and thereby the periodicities present in the pool. Periods longer than 6 nt are difficult to recover with short discrete data, but are present as contributions to fractional periods in the graphs plotting $A_k$ against $p_k$, displayed in Figures 3 and 4 from the main paper. In the absence of specific periodicities, the Fourier amplitude reflects the average pool nucleotide fraction (in percentage) for all periods.

### Supplementary information XIII: Self-complementarity

Self-complementarity related to possible self-binding has been investigated in sequences. This is done by comparing a strand's sequence from the 3′-end to its sequence from the 5′-end and looking for the largest possible complementary overlap. The specific procedure employed in our self-written LabVIEW program then also maximized the area between the two complementary parts of a strand. While secondary structure remains unknown, as well as whether the sequences are actually forming hairpins or just happened to do so in the past or not at all, plotting the average length of self-complementarity against the sequence length permits to discover at which strand lengths self-complementarity becomes a prominent phenomenon. For AT, the fast replicators (long sequences in the middle time point) possess the longest average longest self-complementary regions, Figure XIII.1. The graphs for GC strands do not exhibit longer self-complementary areas than a randomly generated sample with homogeneous average pool nucleotide fraction.

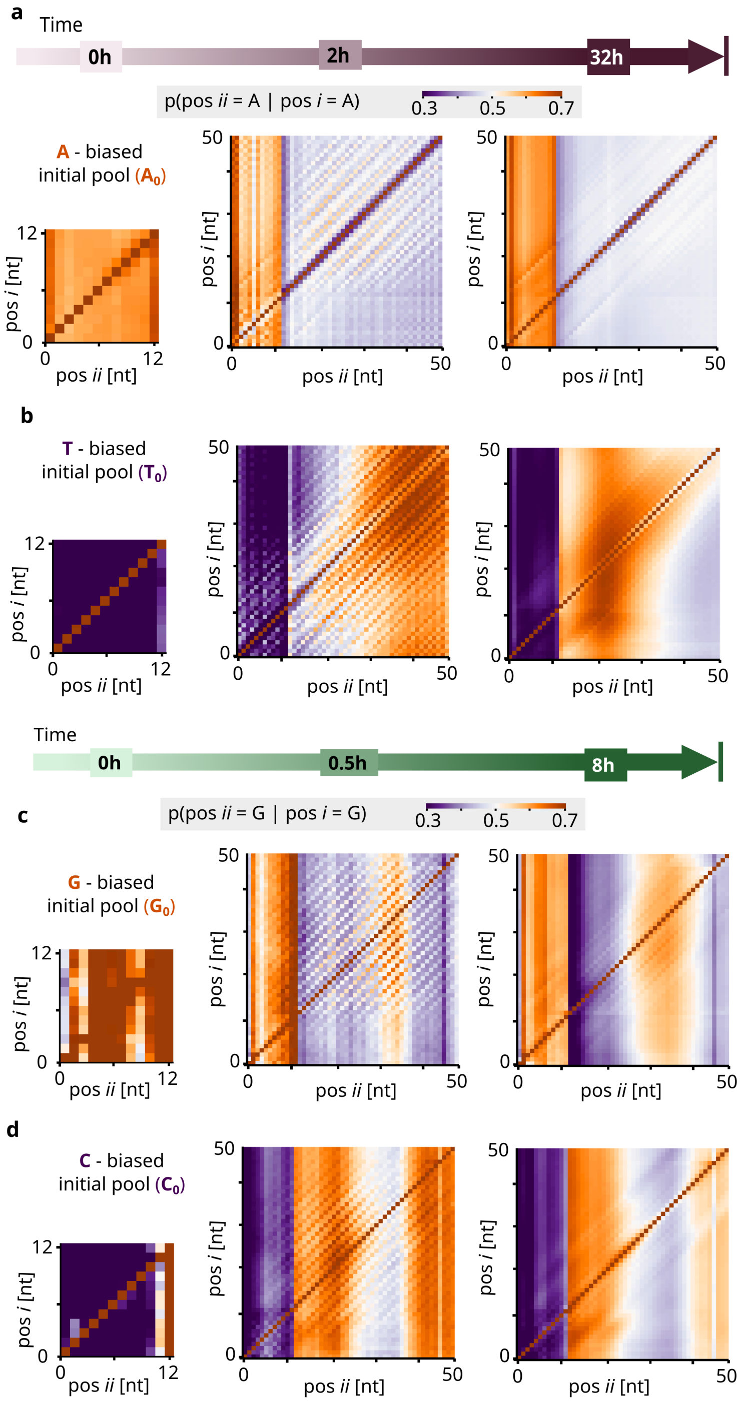


**Figure XII.1:** Conditional probabilities $p(pos\ ii = A/G \mid pos\ i = A/G)$: probability of finding base A (for $A_0$, **a** and $T_0$, **b**) or G (for $G_0$, **c** and $C_0$, **d**) at position *ii*, given A or G at position *i*. The early time points have a mostly homogeneous bias across positions and did not reveal any particular periodic structure. For the intermediate time points, 2h for AT samples and 0.5h for GC samples, diagonal structure indicating periodicity was visible for all the samples. This periodicity, however, was not present in the later time points. These had only vertical regions of bias, corresponding to structures that are present in the average of the pool, as for instance for the initial 12 mer.

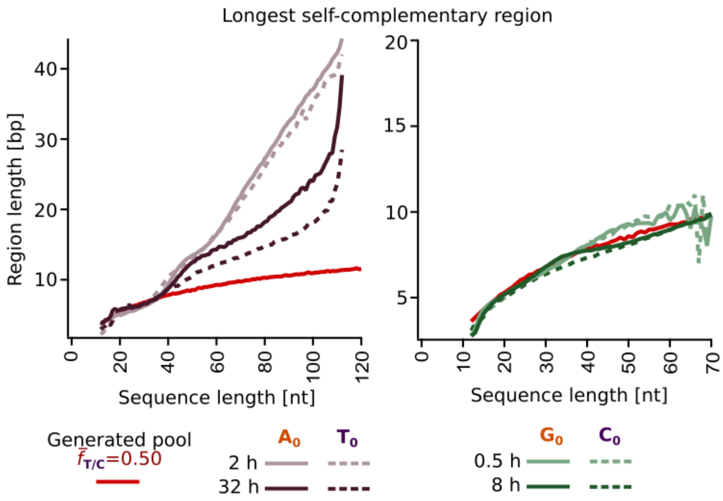


**Figure XIII.1:** Longest self-complementary regions that were found for each sequence, plotted averaged per sequence length. AT samples are show in the left panel and GC samples in the right panel. Additionally, the same analysis was performed for a randomly generated pool with homogeneous length and nucleotide distribution. AT samples showed higher self-complementarity compared to the generated homogeneous pool, particularly for sequences longer than 40 nt. No significant deviations from the randomly generated pool were present in the GC pools.
